# Cooperative herbivory between two important pests of rice

**DOI:** 10.1101/2021.03.10.434708

**Authors:** Qingsong Liu, Xiaoyun Hu, Shuangli Su, Yufa Peng, Gongyin Ye, Yonggen Lou, Ted C. J. Turlings, Yunhe Li

## Abstract

Normally, when different species of herbivorous arthropods feed on the same plant this leads to fitness reducing competition. We found this to be uniquely different for two of Asia’s most destructive rice pests, the brown planthopper and the rice striped stem borer. Both insects directly and indirectly benefit from jointly attacking the same host plant. Double infestation improved plant quality, particularly for the stemborer because the planthopper fully suppresses caterpillar-induced production of proteinase inhibitors. It also drastically reduced the risk of egg parasitism, due to diminished parasitoid attraction. Females of both pests have adapted their oviposition behaviour accordingly. Their strong preference for plants infested by the other species even overrides their avoidance of plants already attacked by conspecifics. This uncovered cooperation between herbivores is telling of the exceptional adaptations resulting from the evolution of plant-insect interactions, and points out mechanistic vulnerabilities that can be targeted to control two major pests.

## Introduction

Species that feed on the same resource are commonly regarded to be competitors ^1-3^. In the great majority of cases, species that share a food resource negatively affect each other’s performance, and the conventional interspecific competition theory is widely recognized for a diverse range of taxonomic groups including plants, birds, reptiles, marine invertebrates, insects and microbes ^1,4,5^. Yet, there are instances of mutually beneficial interactions between species that feed on the same food source. This is mostly known for microbes that can assist other organisms in various ways to help reach, convert and digest food ^6-8^. In very rare cases, higher organisms have also evolved cooperative interactions to be better exploit and share resources, as, for instance, has been shown for predators with complementary hunting tactics ^9-11^.

Amongst arthropods, mutually beneficial interactions have been reported mainly for social insects with food providing partners ^12^. The classic example is the symbiotic relationship between ants and aphids, in which aphids produce sugar-rich honeydew that is collected by the ants, and in exchange ants care for and protect the aphids from predators and parasitoids ^13,14^. In none of these associations the arthropods share the same food sources, and we are not aware of any example of mutually beneficial exploitation of the same resources by arthropods.

The most commonly food sources shared among arthropods are plants, with virtually all plants being attacked by a number of different species. This has so far assumed to always lead to competition ^1,3^. The extent of this competition can vary and the consequences can be highly asymmetrical, but in all reported examples the effects of feeding on the same plant are negative for at least one of the herbivores, independently of whether they are phylogenetically close, have the same mode of feeding, or feed on the same tissues ^1,15-18^. In certain cases, one herbivore species can benefit from the presence of another herbivore species; for instance, by causing physiological changes in the plants that make these plants less toxic and/or more nutritious ^16,17,19-22^ or by masking the (volatile) foraging cues used by natural enemies ^23-27^. To our knowledge, however, there are no known examples of two species of herbivores both consistently benefitting from presence of each other on the same host plants.

It has been postulated that mutually beneficial interactions among species of insect herbivores must exist, but no specific examples have been uncovered yet ^20,28^. It is one thing to demonstrate that two herbivores benefit from jointly feeding on the same plant, it is another to conclude that they actively pursue the plant mediated-benefits. Here we propose such a coevolved partnership between the brown planthopper (BPH), *Nilaparvata lugens* and the rice striped stem borer (SSB), *Chilo suppressalis*, two of the most devastating pests of rice ^29^. Our hypothesis that both insects benefit from attacking the same plant was prompted by our recent finding that BPH can escape parasitism of its eggs by preferentially ovipositing on rice plants that are already infested by SSB ^25^. The apparent reason for this oviposition strategy is that, *Anagrus nilaparvatae*, the most common egg parasitoid of BPH, uses volatiles emitted by BPH-infested plants to locate plants with eggs. Plants that are co-infested by SSB release a different blend of volatiles that is not attractive to the parasitoid ^25^. Previous work also indicates that SSB larvae perform poorly on rice plants that are already infested by conspecifics due to induced plant resistance, and that females show a strong oviposition preference for uninfested rice plants relative to SSB-infested rice plants, in accordance with the ‘mother knows best’ principle ^30^. BPH has been shown to suppress certain defenses in rice ^31,32^. This raises the question whether sharing host plants with BPH can help SSB to counter the direct defenses of rice plants and possibly mitigate the defense responses to SSB infestation, and, if so, whether it also has adapted its oviposition behavior accordingly.

To answer these questions, we tested the performance of SSB larvae on uninfested rice plants, and plants infested either by BPH only, by SSB only, or by both species. We further tested if the oviposition preferences of SSB moths matched the measured performances on the differently pre-infested plants. The results prompted us to investigate the potential molecular and biochemical mechanisms that may be involved, with a focus on protease inhibitors. To also address how BPH may affect the indirect defenses of rice plants against SSB, we further tested the odor preferences of *Trichogramma japonicum*, an important egg parasitoid of SSB, for differently treated plants. With these olfactometer assays and additional cage experiments we tested whether the presence of BPH can also reduce the risk of SSB moth eggs to be parasitized. The combined results support our hypothesis of a co-evolve cooperation between two intimately interacting herbivorous species that feed on the same plants, which exceptionally goes against conventional competition theory of interspecific interactions among phytophagous insects ^1^. The elucidation of the underlying plant-mediated mechanisms that facilitate this cooperation can be the basis for plant breeding strategies to control the two exceedingly important rice pests.

## Results

### Performance of SSB caterpillars on herbivore-infested rice plants

When *C. suppressalis* larvae were allowed to feed for 7 days on rice plants that were either uninfested (control), infested by SSB larvae alone (SSB), BPH nymphs alone (BPH), or both SSB larvae and BPH nymphs (SSB/BPH), the body weight of SSB caterpillars was significantly different among the treatments (*F*_3,165_ = 8.462, *P* < 0.001) (Fig. 1a). The body weight of SSB larvae was significantly lower when fed on plants that had been infested by SSB larvae than on all other plant treatments (all *P* < 0.01). Importantly, there was no difference in larval weight among the other three treatments (uninfested plants, feeding on BPH-infested plants or SSB/BPH-infested plants; *P* > 0.05). These results imply that additional infestation by BPH fully eliminated the negative effects of SSB-infestation on successively feeding conspecifics. And this effect seems to be independent of infestation sequence, as weight gain was only marginally different between SSB larval that were placed on dual infested plants, infested, with SSB treatment occurring either before or after BPH infestation (*P* = 0.051) (Fig. 1b).

**Figure 1.**
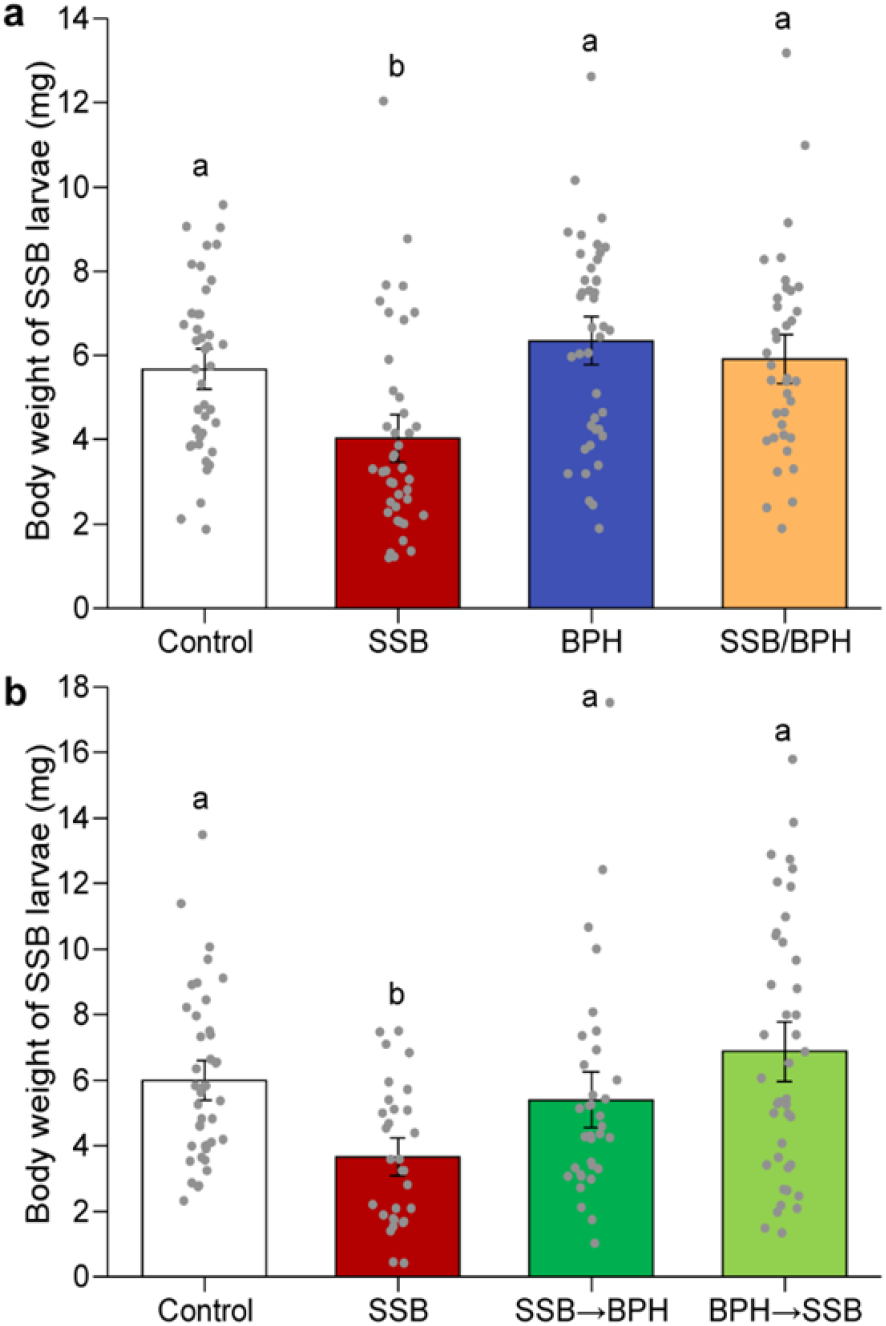
Weight of SSB larvae after seven days of feeding on differentially infested rice plants. **a** Neonates of SSB were individually placed on rice plants that were either uninfested (Control), infested by SSB larvae only (SSB), BPH only (BPH), or both SSB and BPH (SSB/BPH, two species were simultaneously introduced to the plants); **b** Neonates of SSB were individually placed on rice plants that were either uninfested (Control), infested by SSB only, or both SSB and BPH in sequencing order (SSB→ BPH, plants were infested with SSB larvae for 24 h then BPH were added for another 24 h; BPH→ SSB, plants were infested with BPH for 24 h then SSB larvae were added for another 24 h). Bars indicate mean ± SE. Data was analyzed using Two-way analysis of variance followed by LSD post hoc test. Small letters indicate significant differences between treatments (*P* < 0.05) (N = 30–46).

### Oviposition preferences of SSB females

#### Oviposition preference in greenhouse

When given a choice between uninfested and SSB-infested plants, SSB females laid significantly fewer eggs on SSB-infested plants than on uninfested plants (RT-test applied to a GLMM, Poisson distribution error; *P* < 0.001) (Fig. 2a). However, compared to uninfested plants, the females strongly preferred to lay eggs on BPH-infested plants (RT-test applied to a GLMM, Poisson distribution error; *P* = 0.007) (Fig. 2b) or on SSB/BPH-infested plants (RT-test applied to a GLMM, Poisson distribution error; *P* = 0.03) (Fig. 2c). As expected, SSB females also laid significantly more eggs on BPH-infested or SSB/BPH-infested plants (RT-test applied to a GLMM, Poisson distribution error; both *P* < 0.001) relative to SSB-infested plants (Fig. 2e, f), while they laid similar numbers of eggs on BPH-infested and SSB/BPH-infested plants (Fig. 2g).

**Figure 2.**
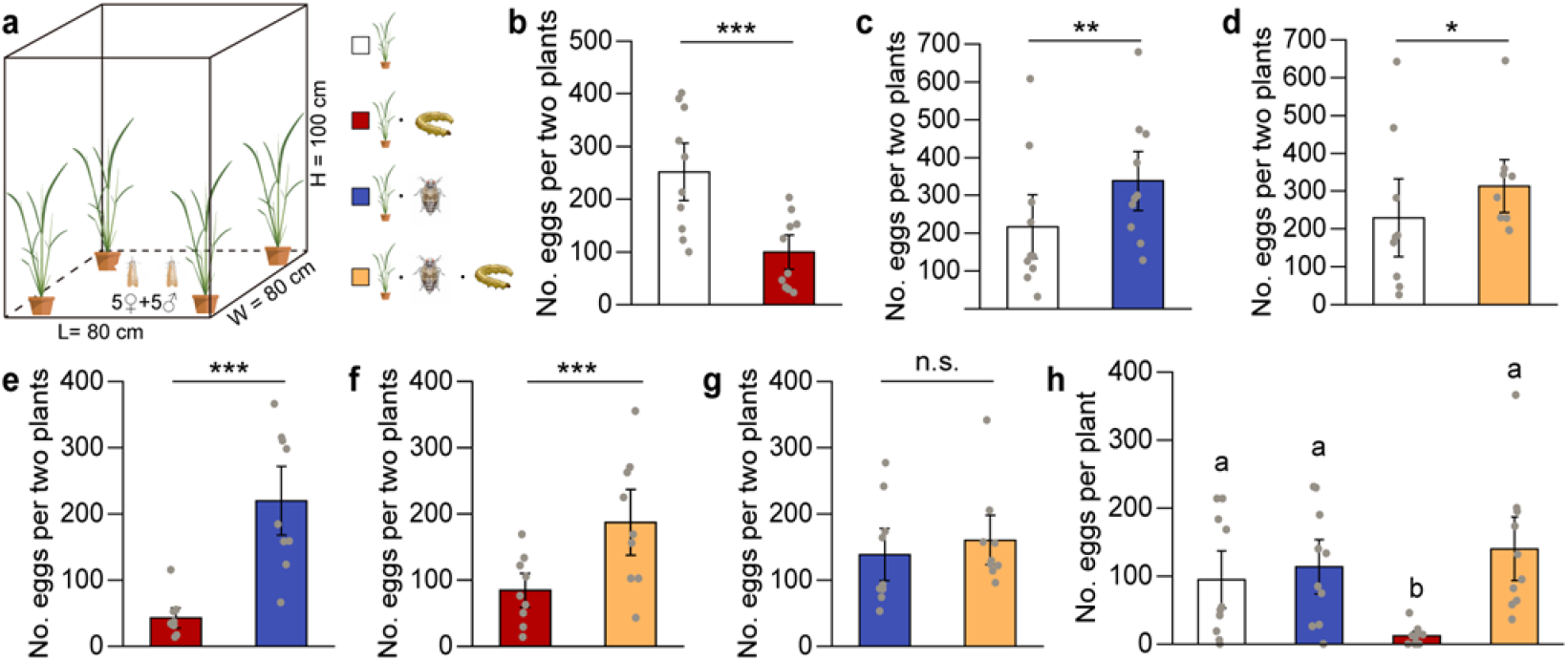
Oviposition preference of SSB female moths in greenhouse experiments. Number of eggs laid by female *C. suppressalis* on rice plants that were either uninfested (Control), infested by SSB larvae only (SSB), BPH only (BPH), or by both SSB and BPH (SSB/BPH). **a** Schematic drawing of the oviposition experiments. **b** Control versus SSB larva-infested plants. **c** Control versus BPH-infested plants. **d** Control versus dually infested plants. **e** SSB larva-infested plants versus BPH-infested plants. **f** SSB larva-infested plants versus dually infested plants. **g** BPH-infested plants versus dually infested plants. **h** SSB female mots were exposed to 4 types of rice plants together. LR tests applied to a GLMM were conducted for the number of eggs (Poisson distribution error). Each bar represents the mean ± SE. Data columns with asterisks (\*\*\**P* < 0.001, \*\**P* < 0.01, \**P* < 0.05, or with small letters (*P* < 0.05) indicate significant differences between treatments; n.s. indicates a non-significant difference (*P* > 0.05) (N = 9–11).

When SSB females were offered the 4 types of rice plants simultaneously, SSB females laid significantly different numbers of eggs among treatments (Fig. 2h; RT-test applied to a GLMM, Poisson distribution error; *P* < 0.001), with the most eggs on plants infested by both SSB and BPH, and slightly less on BPH-infested plants and uninfested plants. Importantly, SSB females laid only very few eggs on SSB-infested plants, significantly less than on the other three treatment (all *P* < 0.001) (Fig. 2h).

#### Oviposition preferences under field conditions

Consistent with the results from the greenhouse experiments, far more eggs were laid on uninfested plants than on SSB-infested plants (RT-test applied to a GLMM, Poisson distribution error; *P* < 0.001) (Fig. 3a). Compared to uninfested rice plants, the females preferred to lay eggs on BPH-infested plants (*P* < 0.001) or SSB/BPH-infested plants (*P* = 0.03) (Fig. 3b, c). When given a choice between SSB-infested plants and SSB/BPH-infested plants, the females preferred to lay eggs on the co-infested plants, as expected (RT-test applied to a GLMM, Poisson distribution error; *P* < 0.001) (Fig. 3d).

**Figure 3.**
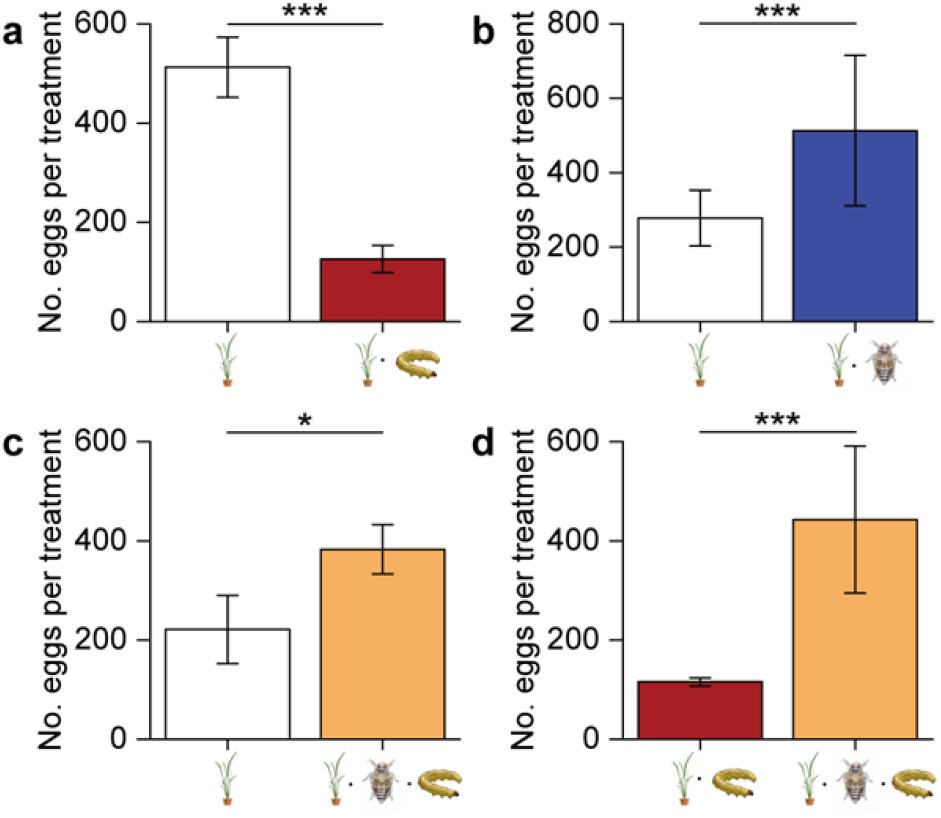
Oviposition preference of SSB female moths in field cage experiments. Number of eggs laid by female *C. suppressalis* on rice plants that were either uninfested (Control), infested by SSB larvae only (SSB), BPH only (BPH), or both SSB and BPH (SSB/BPH). **a** Control versus SSB larva-infested plants. **b** Control versus BPH-infested plants. **c** Control versus dually infested plants. **d** SSB larva-infested plants versus dually infested plants. LR tests applied to a GLMM were conducted for the number of eggs (Poisson distribution error). Each bar represents the mean ± SE. Data columns with asterisks (\*\*\**P* < 0.001, \**P* < 0.05) indicate significant differences between treatments (N = 3–4).

### Rice plant defense responses to herbivore infestation

#### Gene expression changes

RNA-seq analysis was carried out to assess gene expression changes in response to infestation by SSB, BPH or both species. A partial least squares discriminant analysis on all 12 transcriptomic datasets provided a global view of the total gene expression across the four treatments. As shown in Fig. 4a, the first two principal components (PCs) explained 39.5% (PC1), and 13.3% (PC2) of the total variance, respectively. PC1 revealed a clear separation of samples with SSB infestation (SSB and SSB/BPH) from others (BPH and control). And PC2 separated samples with BPH infestation (BPH and SSB/BPH) from others (SSB and control). Taken together, these results suggest that each infestation regime had distinctly different effects on the gene expression of rice plants.

**Figure 4.**
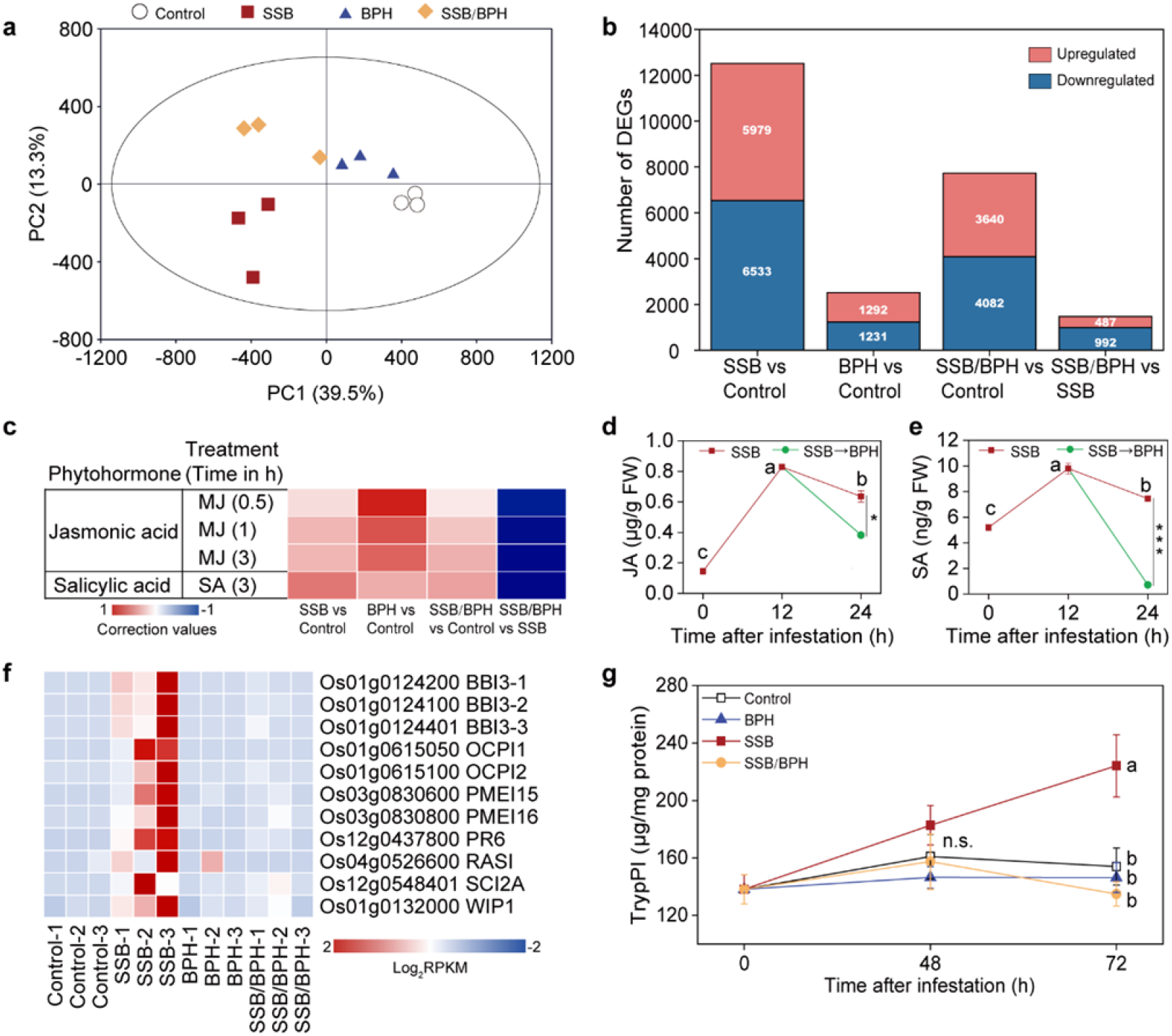
Rice plant responses to herbivore infestation. **a** Partial least squares discriminant analysis (PLS-DA) of all detected genes in rice plants that were either untreated (control), infested by SSB, BPH, or both herbivores (SSB+BPH) for 48 h (three biological replicates per treatment). The percentage of variation of the data explained by principal component 1 (PC1) and PC2 is in parentheses (39.5 and 13.3%, respectively). The score plot displays the grouping pattern according to the first two PCs and the ellipse defines the Hotelling’s T2 confidence interval (95%) for the observations. **b** Differentially expressed genes among differently treated rice plants. **c** Hormonometer analyses for JA and SA signatures based on transcriptomic responses of rice to herbivory. The colors indicate similarity between herbivore infestation and a particular hormone response (blue and red for negative and positive correlations, respectively, see bottom). MJ, methyl jasmonate; SA: salicylic acid. The contents of endogenous JA (**d**) and SA (**e**) in rice plants subjected to SSB infestation or SSB plus BPH in sequence. Letters indicate significant differences among SSB treatment with different time points; asterisks indicate significant differences between SSB and SSB→ BPH treatments (*P* < 0.01; N = 3). Bars indicate mean ± SE. **f** Heatmap of the expression of 10 enzyme inhibitors genes. Log_2_-transformed RPKM values are plotted. BBI 3-1, bowman-birk inhibitor 3-1; BBI 3-2, bowman-birk inhibitor 3-2; BBI3-3, bowman-birk inhibitor 3-3; OCPI1, *Oryza sativa* chymotrypsin inhibitor-like 1; OCPI2, *Oryza sativa* chymotrypsin inhibitor-like 2; PMEI 15, pectin methylesterase inhibitors 15; PMEI 16, pectin methylesterase inhibitors 16; PR6, pathogenesis-related proteins 6; RASI, rice alpha-amylase/subtilisin inhibitor; SCI2A, subtilisin-chymotrypsin inhibitor-2A; WIP1, wound-induced protease inhibitor. FPKM, fragments per kilobase of transcript per million fragments mapped. **g** Time course of the contents of trypPI in rice plants that were either uninfested (Control), infested by SSB, BPH, or both (N=3). Bars indicate mean ± SE. Data was analyzed using one-way ANOVA followed by LSD post hoc test. Letters indicate significant differences between treatments (*P* < 0.05). n.s., not significant (*P* > 0.05).

The gene expression analyses showed that feeding by SSB resulted in the differential expression of 12512 genes (absolute log_2_(fold change) > 0 and *P* < 0.05), of which 6533 genes were up-regulated and 5979 genes down-regulated. Infestation by BPH alone induced a relative weak response of rice plant, that is, 2523 differentially expressed genes including 1292 upregulated genes and 1231 downregulated genes. Co-infestation by the two species induced the upregulation of 3640 genes and the downregulation of 4082 genes (Fig. 4b). Interestingly, compared to SSB-infested plants, 992 genes were downregulated when plants were co-infested with BPH and SSB, and the gene ontology (GO) enrichment analysis showed that these repressed genes were largely associated with defense responses, such as JA-related process, and enzyme inhibitor activity (Supplementary Data 1).

#### JA and SA associated gene expression and their accumulation

A total of 10392 *Arabidopsis* orthologs of rice genes were included in phytohormone-related gene expression signatures (Supplementary Data 2). The results showed that, when compared to uninfested plants, genes associated with JA and SA pathways were generally upregulated in plants infested by SSB larvae, BPH nymphs, or both species (Fig. 4c). However, dual infestation by SSB and BPH resulted in an apparent downregulation of both JA and SA pathways in rice, as when compared to SSB-infested plants. Specifically, four genes involved in JA biosynthesis, *OsLOX9, OsJAR1;2, OsDAD1;3*, and *OsAOC*, were significantly induced upon SSB infestation but were not induced when plants were co-infested by both BPH and SSB (Supplementary Data 3). More surprisingly, two genes involved in JA pathway, *OsLOXL-2* and *OsAOS3*, and nine SA-responsive genes (*OsPR2, OsPR4, OsPR4B, OsPR4C, OsPR4D, OsPR6, OsPR10, OsPR10A*, and *OsPR10B*) were activated by SSB infestation, but were suppressed by dual infestation (Supplementary Data 4).

Consistent with the phytohormone-related gene expression results, SSB infestation induced a significant increase in the levels of both JA (12 hr vs. 0 hr, *P* < 0.001; 24 hr vs 0 hr, *P* < 0.001) and SA (12 hr vs. 0 hr, P < 0.001; 24 hr vs 0 hr, *P* = 0.004; Fig. 4d, e). However, when BPH nymphs were added after 12 hr infestation by SSB alone, the levels of JA and SA in rice plants significantly decreased (JA: *P* = 0.02; SA: *P* < 0.001; Fig. 4d, e).

#### Protease inhibitor-associated gene expression and their accumulation

When determining GO terms that were significantly enriched (*Padj* < 0.05) in the set of 992 downregulated DEGs comparing dual infestation samples and SSB infestation samples, we identified several molecular function terms that were associated with protease inhibitor activity including serine-type endopeptidase inhibitor activity (GO:0004867), enzyme inhibitor activity (GO:0004857), endopeptidase inhibitor activity (GO:0004866), peptidase inhibitor activity (GO:0030414) (Supplementary Data 1). After screening for genes involved in these categories, we found 11 genes related to protease inhibitors that were highly induced by SSB infestation, but were significantly attenuated by the additional infestation of BPH nymphs (Fig. 4f). The expression of nine genes was validated by qRT-PCR, showing similar expression patterns among the four treatments as obtained with RNA-seq (Supplementary Fig. 1), confirming the reliability of the RNA-seq data.

Prompted by the observed changes in protease inhibitor-associated gene expression, we further measured the contents of TPIs in rice plants responding to different herbivore infestations. The results showed a significant increase in TPIs content after 72 hr SSB infestation compared to uninfested plants (*P* = 0.03). This TPIs content increase was not observed for BPH-infested plants or SSB/BPH-infested plants (Fig. 4g).

### Volatile profiles of rice plants and their effects on the oviposition behavior of SSB females

A total of 61 compounds were detected in the headspace of the four plant treatments (Supplementary Data 5). Plants infested by both SSB and BPH emitted the highest amounts of volatiles, followed by plants infested by SSB, plants infested by BPH, whereas control plants emitted the lowest amounts of volatiles. A projection to partial least squares-discriminant analysis (PLS-DA) using the contents of all detected volatiles showed a clear separation among the four treatments (Fig. 5a). The first two significant principle components (PCs) explained 33.8% and 7.44% of the total variance, respectively. Consistent with gene expression data, the first PC showed a clear separation between plant infested with SSB (SSB and SSB/BPH) from others (BPH and control), and the second PC separated BPH-infested samples and SSB-infested samples (BPH and SSB) from others (SSB/BPH and control).

**Figure 5.**
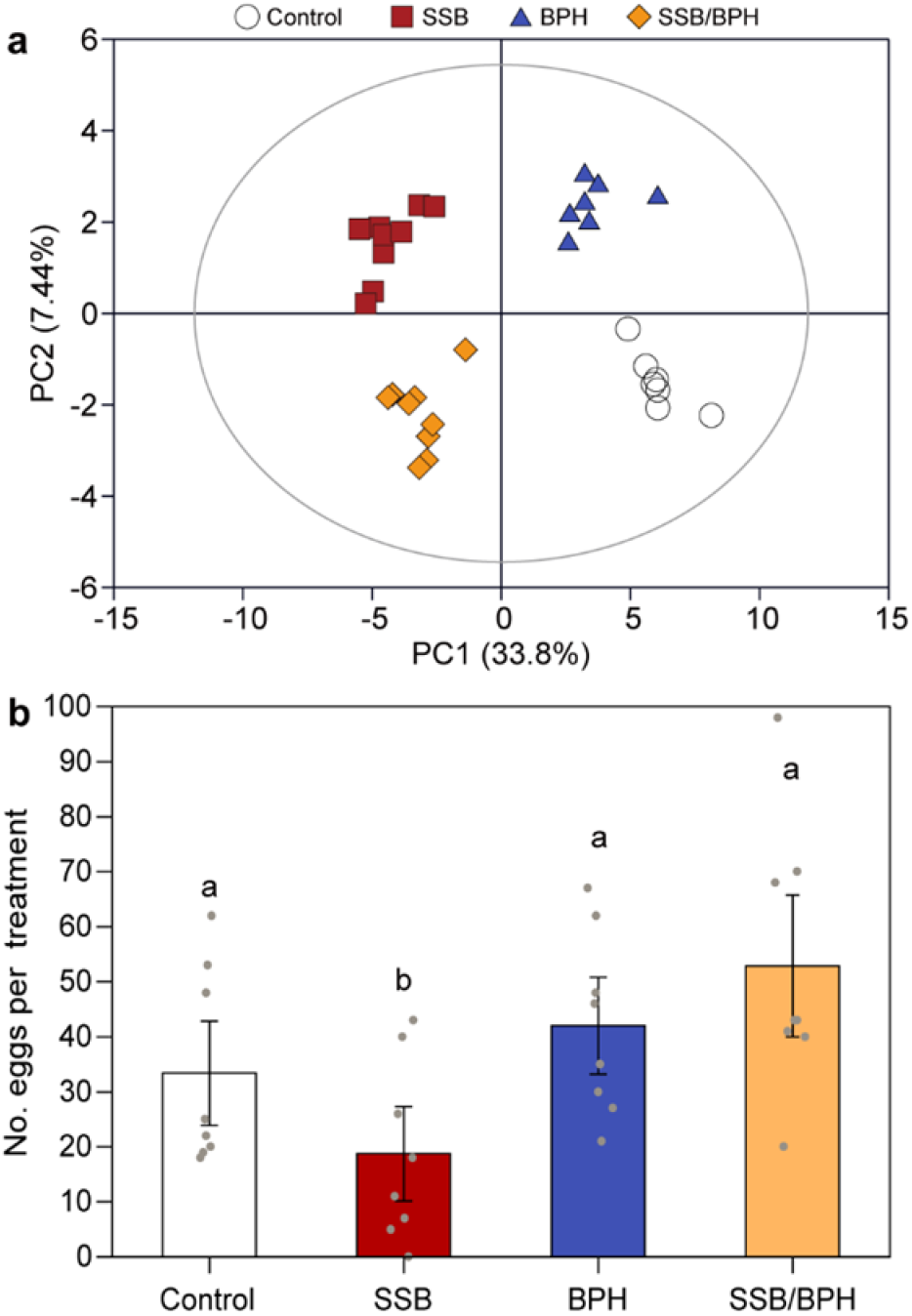
Volatiles released by rice plants and effect on oviposition behavior of *C. suppressalis* female. **a** Projection to latent structures-discriminant analysis (PLS-DA) of volatile emissions produced by rice plants that were either untreated (Control), infested by SSB, BPH, or both herbivores (SSB/BPH) for 48 h. The score plot display the grouping pattern according to the first two components and the ellipse defines the Hotelling’s T2 confidence interval (95%) for the observations. **b** Number of eggs laid by female *C. suppressalis* on filter paper that had been treated with volatiles collected from 4 types of differently treated rice plants. LR tests applied to a GLMM were conducted for the number of eggs (Poisson distribution error). Each bar represents the mean ± SE. Letters above bars indicate significant differences between treatments (RT-test applied to a GLM, Poisson distribution error; *P* < 0.05; N = 8).

Using volatiles that had been collected from plants subjected to the four types of treatments as odor sources (applied on filter paper strips), we tested the oviposition preference of SSB females. They differently distributed their eggs among the four treatments (Fig. 5b, RT-test applied to a GLMM, Poisson distribution error; *P* < 0.001), with highest number of eggs observed on or near filter paper that had been treated with volatiles collected from SSB/BPH-infested plants, which was statistically similar to the numbers of eggs laid on BPH-infested plants and uninfested plants. However, SSB females laid significantly lower numbers of eggs on filter paper treated with volatiles collected from SSB-infested plants compared to any of the other treatment (all *P* < 0.001) (Fig. 5b).

### Responses of *T. japonicun* wasps to herbivore-infested rice plants

In a dual-choice assay, the *T. japonicun* wasps showed a strong preference for plants infested by SSB eggs (*P* = 0.004) over uninfested intact plants (Fig. 6a). When offered plants carrying SSB eggs in both sides of the olfactometer, the wasps exhibited a significant preference for plants that were additionally infested by SSB larvae relative to plants infested by SSB eggs alone (*P* = 0.002). In contrast, the wasps were significantly more attracted to plants with eggs alone than plants that were additional infested by BPH nymphs (*P* < 0.001) or both SSB larvae and BPH nymphs (*P* = 0.001). We further tested the preferences of wasps that were offered plants with combinations of herbivore and egg infestation. The results (Fig. 6a) showed that *T. japonicun* females significantly preferred to rice plants with SSB and eggs over plants coinfested by BPH and eggs (*P* < 0.001) or plants with SSB, BPH, and eggs (*P* = 0.01). Finally, as expected, the wasps showed a significant preference for the odor of plants with the combination of SSB, BPH, and eggs over the odor of plants with BPH and eggs (*P* = 0.009). These results imply that additional infestation by BPH resulted in the repellence of *T. japonicun* wasps.

**Figure 6.**
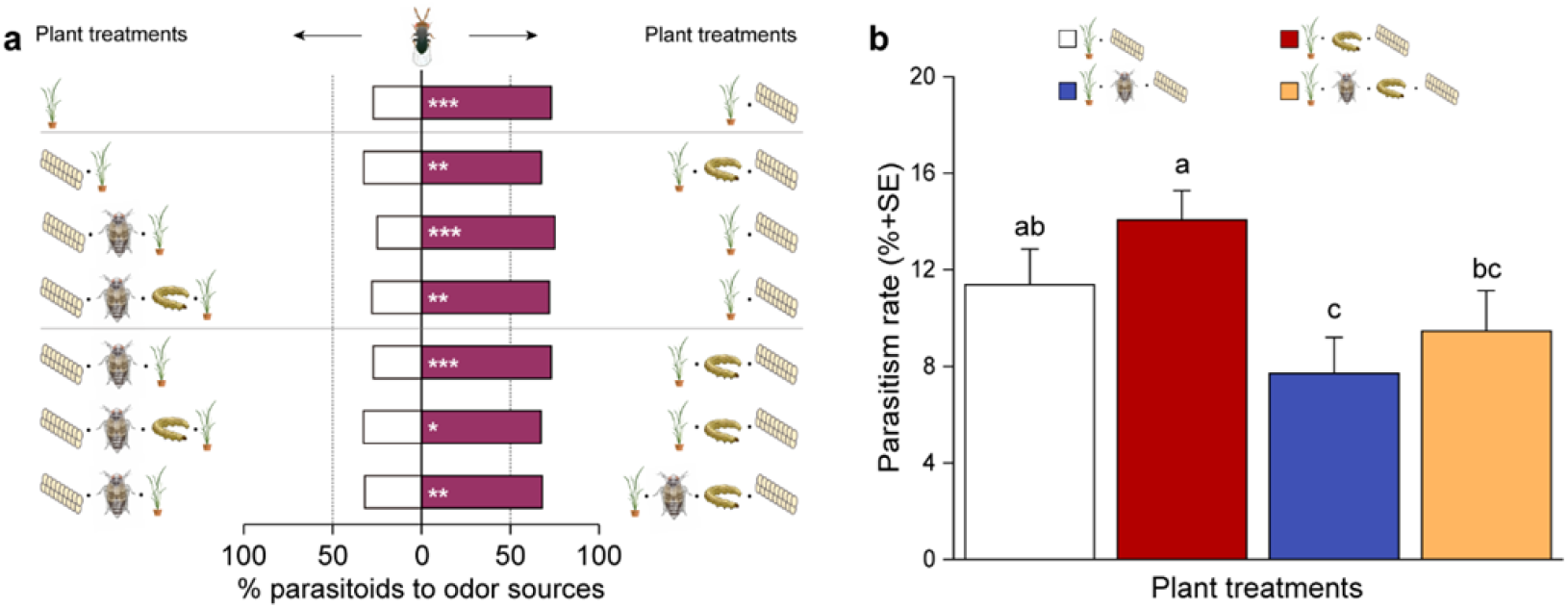
Preferences of females of the egg parasioid *Trichogramma japonicum*. **a** Choice of *T. japonicun* wasps when offered the odor of differently treated plants in a Y-tube olfactometer. Bars represent the percentages of wasps choosing either of the odor sources. Asterisks indicate significant differences from a 50:50 distribution (binomial test: * *P* <0.05, ** *P* < 0.01, *** *P* < 0.001; N =58-80). **b** Parasitism rates of SSB eggs by *T. japonicum* in the greenhouse experiments. The proportions data was subjected to an arcsin square-root transformed before analyses. Each bar represents the mean ± SE. Letters above bars indicate significant differences between treatments (likelihood ratio test applied to a generalized lineal model; *P* < 0.05; N = 12).

### Parasitism rates of SSB eggs by *T. japonicun* wasps

In the greenhouse cages with the four plant treatments, the rate of parasitism of SSB eggs by *T. japonicum* wasps were highest on plants infested with SSB larvae only (Fig. 6b), which was significantly higher than on plants infested with BPH only (*P* < 0.001) or plants infested with both herbivore species (*P* = 0.016). The lowest parasitism rate of SSB eggs was observed on plants infested by BPH only. Although parasitism was lower on plants infested by both species, this was not significant different from parasitism on control plants (*P* = 0.28). The parasitism rates of SSB eggs in the cages nicely reflected the trends of responses of the parasitoids in the olfactometer (Fig. 6b).

## Discussion

Defining *cooperation* as “any interaction in which an actor confers a fitness benefit to another individual and receives an (inclusive) fitness benefit in return”^33^, we may conclude that the observed mutually beneficial interactions between SSB and BPH indeed represent a unique example of cooperation between two herbivores. We found that not only do the herbivores directly (mitigated plant toxicity) and indirectly (reduced exposure to parasitoids) benefit from jointly attacking plant, we also found that both have adapted their host plant selection and oviposition behavior to optimize the benefits that they derive from each other.

Our previous work shows that BPH prefers to feed and lay eggs on SSB-infested rice plants, which are more nutritious and on which their eggs escape parasitism ^25,34^. The current study demonstrates similar, even stronger benefits for SSB when infesting plants that are already under attack by BPH. The feeding assays show that SSB infestation induces rice defense responses that cause significantly reduced fitness in SSB larvae that subsequently feed on the same plants (Fig. 1). Remarkably, additional infestation by BPH completely neutralized the negative effects of SSB defense induction; SSB caterpillars placed on plants already infested by conspecifics grew considerably better on plants that were also attacked by BPH, independent of order of infestation. On BPH-infested plants the caterpillars performed just as well as on previously uninfested control plant (Fig. 1)

RNA-seq and biochemical analyses showed that infestation by BPH suppresses a broad spectrum of defense related genes (Fig. 4f, Supplementary Data 1), with one of the main consequences being a significant reduction in SSB-induced production of proteinase inhibitors (Fig. 4g). These are key defensive compounds of rice plants that are particularly effective against chewing herbivores, including SSB ^35,36^. We also found that co-infestation with BPH suppresses the expression of SA- and JA-associated genes that are normally upregulated by SSB infestation (Fig. 4c, Supplementary Data 4). This suppression led to reduced levels of JA and SA in the plants (Fig. 4d, e). The JA signal transduction pathway is responsible for the production of TPIs in rice plant, and it is known that SSB performs better on JA-deficient mutant rice lines mainly due to reduced TPIs levels ^36,37^. Collectively, this insight explains why BPH feeding neutralizes the negative effects of SSB defense induction.

Insect attack typically induces defense responses in plants ^38-41^, but it is also increasingly evident that various insect herbivores have the ability to, at least partially, suppress plant defenses by interfering with JA or SA biosynthesis and that this can enhance their own performance and that of conspecifics ^28,42-45^. This can also benefit certain other species that feed on the same plants. For example, the silverleaf whitefly (*Bemisia tabaci*) activates SA signaling and represses JA-regulated defenses, leading to enhanced nymphal development of this insect ^28^, which also benefits spider mites ^26^. None of these studies appear to show reciprocal benefits for species feeding on the same plant.

Consistent with ‘mother knows best’ hypothesis^46^, SSB females avoid to lay eggs on rice plants that are already infested by conspecifics, thus ensuring that their offspring evade the negative effects that SSB-induced defenses ^30^. Here we show that SSB females have adapted their oviposition behavior to preferentially oviposit on BPH-infested rice plants, independent of whether SSB larvae are already present or not, as compared to healthy plants (Figs. 2 and 3). By doing so, they benefit from the BPH-mediated suppression of rice defense responses (Fig. 1a). However, the performance of SSB larvae was not any better on rice plants that were already co-infested by SSB plus BPH compared to their performance on healthy plants. So why did SSB females prefer to lay eggs on dual-infested plants rather than on healthy plants (Fig. 2d)? The experiments with the egg parasitoid *T. japonicun* provide a plausible explanation, as they showed that the presence of BPH significantly reduced the attractiveness of rice plants to the wasp (Fig. 6a). The cage experiment confirmed that the presence of BPH decreased the risk for SSB eggs to be parasitized, implying that the oviposition strategy of SSB females is highly adaptive (Fig. 6b).

It appears that certain well-adapted herbivores can also manipulate the emission of volatiles of their host plants ^47^. In most such cases, herbivore infestation suppresses certain key volatile compounds and thereby possibly reduce the attractiveness of plants to certain natural enemies ^26,27,42,48,49^. Other herbivores can benefit from this as well. Simultaneous feeding by slugs, for instance, suppress caterpillar-induced volatiles in cabbage plants, thereby reducing the attractiveness of the plants to parasitoids ^27^. Similarly, when spider mite-infested Lima bean plants are also infested by whiteflies this leads to a reduced emission of the volatile (E)-β-ocimene compared to plants infested by spider mites only, resulting in a reduced attraction of predatory mites ^26^. In other cases, double infestation may actually lead to higher quantities of volatiles being emitted, but the blend is altered in a way that it is no longer attractive to parasitoids ^23^. This is also the case for our study system. We previously showed that BPH preferentially oviposits on SSB-infested rice plants, thereby avoiding the attraction of the egg parasitoid *Anagrus nilaparvatae* ^25^. These various examples confirm that insect herbivores not only can evolve the ability to use volatiles to identify host plants of better nutritional quality, but also plants where their offspring can escape natural enemies ^50^.

Our combined results, including the additional tests showing that SSB infestation of rice plants significantly increases the performance of BPH nymphs (Supplementary Fig.2) strongly supported the conclusion of mutually beneficial interaction between SSB and BPH. This goes against the ingrained notion of competition between phytophagous insects that share a common host plant, and how this competition shapes insect assemblages ^1,15,17,51-53^. The resource-based competition theory has been challenged before, but the examples involve specific asymmetric beneficial plant-mediated interactions, meaning that only one of the herbivore species benefits from the presence of another ^20,21,54-58^. In these cases, the benefit is never reciprocal, nor do they seem to represent tightly coevolved interactions with specific behavioral adaptations as found for BPH and SSB in the current study and certain vertebrates ^9-11^.

The interaction between SSB and BPH reported here seems to represent a highly evolved collaboration to cope with and exploit the direct and indirect defense responses of rice plants. The two species share the same host plants and have a similar spatial and temporal distribution throughout Asia’s rice paddy area (^25^; https://www.cabi.org/isc/). Why has their interaction evolved into collaboration rather than competition? It is likely because of their differential feeding strategies. Although the two insects occur side-by-side on rice plants, SSB is a stemborer insect that feeds inside the rice plants, while BPH is a phloem-sucking insect that feeds at the surface on leaf blades and leaf sheaths ^25,59^. Hence, there is usually no direct physical interaction between the two species. During the coevolutionary arms race between herbivores and host plants, both sides may evolve multiple defense mechanisms against the other. We speculate that the cooperative relationship between SSB and BPH may the result of two opposing coevolutionary arms races that in combination benefit both herbivores.

Although the mutually beneficial interaction between the stemborer and planthopper bears no resemblance to any of the known interactions between other herbivore species attacking a same plant ^1,15^, the reported type of cooperation is unlikely to be unique. We postulate that agricultural pests are especially prone to rapid evolutionary changes that allow such cooperation to emerge. As pest populations build up in vast monocultures their main challenges revolve around coping with plant defenses and avoiding and resisting their specific natural enemies, whereas finding host plants is no longer a challenge. In such scenarios, different insect species will encounter each other frequently. Unlike the cultivated plants, the insects are subject to natural selection and can evolve traits to jointly overcome plant defenses, without the cultivated plants being able to coevolve to resist these traits. The plants are at the mercy of human selection, which is focused at traits that favor yield and nutritional value, often at the cost of reduced resistance against pests and diseases ^60,61^. Yet, as we discover and unravel the intricate adaptations in the insects we can start steering this human selection in favor of potent pest resistance traits. For the specific example uncovered in our study, interfering with the ability of BPH to suppress the biosynthesis of proteinase inhibitors could be a highly effective and sustainable strategy to control two of rice’s most common and most harmful pests.

In summary, the current study reveals a highly adaptive, mutually beneficial relationship between rice planthoppers and stem borers that is mediated by opposing rice plant defense responses. The findings represent a unique example of a cooperative interaction that challenges traditional interspecific competition theory. The two insect species take advantage of the rice defense responses induced by each other in a manner that suggests that together they are the tentative winners in the arms race with rice plants. The results are also illustrative of the complexity and intricate dynamics of the interaction between plants and insects, and challenge the conventional paradigms of interspecific competition. Future work should further unravel more details about the molecular mechanisms underlying the insect-controlled interactions, which might lead the development of rice varieties that disrupt the cooperative interaction as potential strategy to control the two pests.

## Methods

### Plants and insects

Rice plants (*Oryza sativa*, cultivar Minghui63) were grown in a greenhouse at 27 ± 3 °C with 75 ± 10% RH (relative humidity) and a photoperiod of 16:8 hr L:D (light:dark). The cultivation of rice plants followed the same procedure as described previously ^25^. Plants were used for experiments when they were at the tillering stage, which occurred about 44–49 days after sowing.

*C. suppressalis* larvae were reared on an artificial diet as described ^62^. Ten percent honey water solution was provided to supply nutrition for the adults. *N. lugens* were maintained on a BPH-susceptible rice variety Taichung Native 1 (TN1) ^34^. *T. japonicun* were obtained from Keyun Industry Co., Ltd (Jiyuan, China). Newly emerged adult wasps were maintained in glass tubes (3.5 cm diameter, 20 cm height) and supplied with 10% honey water solution as a food source and were maintained for at least 6 hr to ensure freely mating, before females were used for the following experiments. All three species were maintained in climatic chambers at 27 ± 1°C, 75 ± 5% RH, and a photoperiod of 16:8 hr L:D.

### Performance of caterpillars on insect-infested rice plants

Multiple types of rice plants were prepared: i) uninfested plants, meaning that potted rice plants remained intact without insect infestation; (ii) SSB-infested plants, each potted rice plant was artificially infested with one 3^rd^ instar SSB larva that had been starved for > 3 hr for 48 hr; iii) BPH-infested plants, each potted rice plant was artificially infested with a mix of fifteen 3^rd^ and 4^th^ instars BPH nymphs for 48 hr; iv) SSB/BPH-infested plants, each potted rice plant was simultaneously infested with one SSB larva and 15 BPH nymphs for 48 hr; (v) SSB→ BPH-infested plants, each potted rice plant was artificially infested with one SSB larvae alone for the first 24 hr, then 15 BPH nymphs were additionally introduced for another 24 hr; vi) BPH→ SSB-infested plants, namely each potted rice plant was artificially infested with 15 BPH nymphs for the first 24 hr, then one SSB larvae were additionally introduced for another 24 hr. Plant treatments were conducted as described in detail in our previous study ^25^. During herbivory treatment, the uninfested plants were placed in a separate room to avoid possible volatile-mediated interference. During the subsequent bioassays, both SSB caterpillar and BPH nymphs remained in or on the rice plants.

Two bioassays were conducted to test the performance of *C. suppressalis* larvae feeding on differently treated rice plants. The first bioassay included the plants treatments i, ii, iii, and vi, and the second bioassay included the plants treatments i, ii, v and vi. Three 2-day-old larvae of *C. suppressalis* were gently introduced onto the middle stem of each rice plant using a soft brush. The infested rice plants were then placed in climatic chambers at 27 ± 1°C, 75 ± 5% relative humidity, and a photoperiod of 16:8 hr L:D. The *C. suppressalis* larvae were retrieved from the rice plants after 7 days feeding, and they were weighed on a precision balance (CPA2250, Sartorius AG, Germany; readability = 0.01 mg). The mean weight of the three caterpillars on each plant was considered as one biological replicate. The experiment was repeated four times using different batches of plants and herbivores, resulting in a total of 30–46 biological replicates for each treatment.

### Oviposition-preferences of *C. suppressalis* females choosing among differently infested rice plants

#### Greenhouse experiment

In the greenhouse, seven choice tests were conducted with *C. suppressalis* females including i) SSB-infested plants versus uninfested plants; ii) BPH-infested plants versus uninfested plants; iii) SSB/BPH-infested plants versus uninfested plants; iv) SSB-infested plants versus BPH-infested plants; v) SSB-infested plants versus SSB/BPH-infested plants; vi) BPH-infested plants versus SSB/BPH-infested plants; and vii) the test in which *C. suppressalis* females were exposed to all four types of rice plants. The experiments were performed as described in detail by ^30^. In brief, four potted plants were positioned in the 4 corners of a cage (80 cm x 80 cm x 100 cm) made of 80-mesh nylon nets for each test. For paired comparisons, two potted plants belonging to the same treatment were placed in opposite corners of each age, and in the test with 4 types of rice plants, each type of plant was positioned in one of the four corners of each cage. Five pairs of freshly emerged moths (less than 1 day) were released in each cage, and a clean Petri dish (9 cm diameter) containing a cotton ball soaked with a 10% honey solution was placed in the center of the cage as food source. After 72 h, the number of individual eggs on each plant were determined. The experiment was conducted in a greenhouse at 27 ± 3°C, 65 ± 10% RH, and a photoperiod of 16:8 hr L:D. Each choice test was repeated with 9–11 times (replicates).

#### Field cage experiment

The oviposition preference of SSB females was further assessed in a field near Langfang City (39.58° N, 116.48° E), China. Four choice tests were conducted: i) SSB-infested plants versus uninfested plants; ii) BPH-infested plants versus uninfested plants; iii) SSB/BPH-infested plants versus uninfested plants; and iv) SSB/BPH-infested plants versus SSB-infested plants. The treated rice plants were prepared as described above and were transplanted into experimental plots (1.5 m × 1.5 m). For each pairwise comparison, six plots of rice plants were covered with a screened cage (8 m × 5 m × 2.5 m) made of 80-mesh nylon net to prevent moths from entering or escaping. Each of the six plots contained 9 rice plants of a particular treatment, with 3 plots per cage representing the same treatment. Plots were separated by a 1-m buffer and they were alternately distributed in a 3 × 2 grid arrangement in each cage (Supplementary Fig. 1). Approximately 50 mating pairs of newly emerged *C. suppressalis* adults (< 24 hr) were released into each cage. After 72 hr, the number of individual eggs on each plant were determined. The total number of eggs of three plots in each cage was regarded as one replicate, 3–4 replicates were conducted for each pairwise comparisons.

### Rice plant response to herbivore infestation

#### RNA-seq and data analysis

To explore the molecular mechanisms underlying the rice plant-mediated interaction between BPH and SSB, gene expression changes in rice response to infestation by SSB, BPH or both were analyzed by RNA-seq. The rice plants, uninfested (control) or infested, were prepared as described above. After 48h, the stems of the plants were harvested and frozen in liquid nitrogen. Samples from 5 individual plants of the same treatment were pooled together as one biological replicate, and three replicates were collected for each treatment.

RNA-seq analyses were performed as described previously ^63^. In brief, total RNA was extracted using the TRIzol reagent (Invitrogen, Carlsbad, CA, USA) and treated with RNase-free DNase I (NEB, Ipswich, MA, USA) to remove any genomic DNA. Library preparation and RNA-seq were performed by Novogene (Beijing, China) using an Illumina Hiseq 4000 system, resulting in ∼45–55 million raw reads per sample. Raw reads were subjected to quality checking and trimming to remove adapters, poly-N sequences, and low-quality bases (Phred quality score Q < 20). The yield clean data of each sample were aligned to the rice reference genome IRGSP-1.0 (https://rapdb.dna.affrc.go.jp) using HISAT2 (v2.09) ^64^, and the number of reads mapped to each gene was counted with featureCounts (v1.5.0-p3) ^65^. The expression level of each gene was calculated as FPKM (fragments per kilobase of transcript per million per million fragments mapped) according to an established protocol ^66^. Expression differentiation analyses were conducted DESeq2 R package (v. 1.18.0) ^67^. Genes with absolute value of log_2_(fold change) > 0 and *P*-value < 0.05 were defined as differentially expressed genes (DEGs). The enriched functions of DEGs in RNA-seq data sets were annotated with the Gene Ontology (GO) function using the clusterProfiler R package ^68^, and GO terms with Benjamini– Hochberg false discovery rate (FDR) adjusted *P*-value (*Padj*) < 0.05 were considered significantly enriched. The transcriptional signatures of hormonal responses of rice plant to herbivory relative to gene expression in *Arabidopsis* induced by diverse phytohormones was analyzed using Hormonometer program ^69^. Since trypsin protease inhibitors (TPIs) serve as indicators of induced resistance in rice plants, especially against chewing herbivores such as SSB ^35,36^, the analyses focused on expression profiles of TPIs-related genes among the four plant treatments. The expression of nine selected TPIs genes were validated by quantitative real-time PCR (qRT-PCR) analyses as previously described ^70^. qRT-PCR was conducted on a Bio-RadCFX96 Touch Real-time PCR Detection System instrument (Bio-Rad, Hercules, CA, USA) using TransStart^®^ Top Green qPCR SuperMix (TransGen Biotech, Beijing, China). The rice *ubiquitin 5* gene was used as the internal standard to normalize the variations in gene expression. The primers used are listed in Supplementary Table 1.

#### Quantification of endogenous jasmonic and salicylic acid

Our RNA-seq results suggested that additional infestation by BPH significantly suppressed the expressions of genes related to the defense hormones jasmonic acid (JA) and salicylic acid (SA). Both types of genes are highly upregulated in response to SSB infestation ^71^. To confirm this, we quantified the JA and SA levels in rice plants with two treatments: i) rice plants that were infested with one third-instar SSB larva alone for 24 hr; ii) rice plants that were first infested with one third-instar SSB larva for 12 hr and then also with 15 BPH female adults for another 12 hr. Rice stems were harvested at three time points: 0 hr (uninfested control plants), as well as 12 hr and 24 hr after infestation. For each treatment, stems from 5 individual plants were harvested and pooled together as one biological replicate, and three replicates were collected for each time point.

Endogenous measurements of JA and SA were performed by the plant hormone platform at the Institute of Genetics and Developmental Biology, Chinese Academy of Sciences as previously described ^72^.

#### Quantification of TPIs

We further measured the accumulation of TPIs in rice plants subjected to insect infestation. These experiments were prompted by the RNA-seq results indicating that the upregulation of TPIs-related genes in response to SSB infestation is significantly suppressed after co-infestation with BPH. The same plant treatments were included as used for RNA-seq but with new batches of plants. Samples were collected at 48 hr and 72 hr of insect infestation. Intact rice plants that served as controls were also sampled at the same time points. Samples from five individual plants were pooled together as one biological replicate, and three replicates were collected for each treatment. All samples were immediately frozen in liquid nitrogen and stored at −80°C until further analyses.

TPIs contents was determined using enzyme linked immunosorbent assay (ELISA) kits (J&L Biological, Shanghai, China). The stem samples were ground into a fine power in liquid nitrogen using a mortar and pestle, and each sample (0.1 g) was homogenized in 0.01 M PBS (Phosphate Buffered Saline) buffer (pH = 7.4) (Sigma-Aldrich, St. Louis, MO, USA) with a sample–PBS proportion of 1:9 (1 g plant sample/9 ml of PBS). Samples were centrifuged at 4000 *g* for 15 min at 4°C and the supernatant was collected. The ELISA experiments were performed following the protocols provided with the kits. The optical density values were recorded at 450 nm using a microplate spectrophotometer (PowerWave XS2, BioTek, Winooski, VT, USA). The protein concentrations in plant samples were measured using a bicinchoninic acid (BCA) protein assay kit (Aidlab Biotechnologies Co., Ltd., Beijing, China) according to the manufacturer’s instructions. The amount of protease inhibitor was calculated based on a standard curve, and results were expressed as µg protease inhibitor per mg protein.

### Effect of insect-induced volatiles on the oviposition behavior of SSB moths

#### Collection and analysis of rice plant volatiles

Individual rice plants were either uninfested or infested with SSB larvae alone, BPH nymphs alone, or both species simultaneously for 48 hr using the method described above. The emitted volatiles were trapped for 3 hr (21:00–24:00, the time period that SSB lay their eggs), and then analyzed and identified as described ^25^. The compounds were quantified as a percentage of peak areas relative to the internal standard (nonyl acetate) per 3 hr of trapping for one plant. For each treatment, collections and analyses were repeated 7–9 times.

#### *Odor preferences of SSB* females

The response of SSB females to volatiles released from differently treated rice plants were investigated to better understand the mechanism underlying the moth’s oviposition preferences. The total volatiles emitted from uninfested plants, SSB-infested plants, BPH-infested plants and SSB/BPH-infested plants were collected for this experiment. Plant treatments and volatiles collections were the same as described above but without the addition of the internal standard. The collected volatiles were diluted in paraffin oil (purity 99%; Sigma-Aldrich, St. Louis, MO, USA) at 1:4 (v/v) and were stored at −80 °C before use.

One ml of each of the four types of volatile solutions were separately pipetted on the center of a filter paper strip (4 cm × 21 cm), which were then hung from the four corners of a cage (45 cm × 45 cm × 45 cm) made of 80-mesh nylon net. Five pairs of freshly emerged SSB moths (< 24 hr) were released in each cage. After 72 hr, the number of eggs deposited on the filter paper strips and the surface of the nylon nets near each paper strip were determined. This oviposition choice test was repeated 8 times.

### Response of the egg parasitoid *T. japonicun* to herbivore-infested rice plants

Multiple types of herbivore-infested rice plants were prepared: i) uninfested plants (control); ii) SSB-infested plants; iii) BPH-infested plants; iv) SSB/BPH-infested plants; v) plants infested with SSB eggs (referred to as egg-infested plants); vi) plant infested with SSB larvae and their eggs (referred to as SSB/egg-infested plants); vii) plants infested with BPH nymphs and SSB eggs (BPH/egg-infested plants); and viii) plants infested with both SSB larvae, BPH nymphs and SSB eggs (referred to as SSB/BPH/egg-infested plants). To prepare these treatments, plants were first artificially infested with herbivores for 48 hr as described above, then some of them were subjected to SSB eggs deposition. For that, two potted rice plants of the same type were placed in a cage (45 cm × 45 cm × 45 cm) made of 80-mesh nylon nets, then 30 pairs of freshly emerged moths (< 24 hr) were released in each cage to mate and lay eggs. After 24 hr, the plants were removed from the cage and those that carried 200-250 eggs were used as odor sources. During the period of egg deposition and the subsequent olfactometer experiments with the parasitoid, all insects remained in or on the rice plants.

To test the behavioral responses of *T. japonicum* to differently treated rice plants, they were offered the following pairs of odor sources: i) uninfested plants versus egg-infested plants; ii) uninfested plants versus SSB-infested plants; iii) uninfested plants versus BPH-infested plants; iv) egg-infested plants versus SSB/egg-infested plants; v) egg-infested plants versus BPH/eggs infested plants; vi) SSB/egg-infested plants versus BPH/egg-infested plants; vii) egg-infested plants versus SSB/BPH/egg-infested plants; viii) SSB/egg-infested plants versus SSB/BPH/egg-infested plants; and ix) BPH/egg-infested plants versus SSB/BPH/egg-infested plants.

Responses of *T. japonicun* females to these odor sources were investigate in a Y-tube olfactometer as described ^25^. Newly emerged adult wasps were maintained in glass tubes (3.5 cm diameter, 20 cm height) for at least 6 hr to ensure that they would mate, before females were used for the experiments. Two rice plants of the same treatment were enclosed in a glass bottle and used as one odor source, and each pair of odor sources was replaced after ten parasitic wasps were tested. For each treatment, a total of 64–88 female wasps were tested. The experiments were conducted between 10:00 and 16:00 on several consecutive days.

### Parasitism rates of *C. suppressalis* eggs by *T. japonicun* wasps

In a cage experiment, we further tested if the differences in parasitoid attraction observed in the olfactometer for the differentially infested plants can result in differences in parasitism rates of SSB eggs under realistic conditions. The following herbivore-treated plants were prepared as described above: SSB eggs on uninfested, SSB-infested, BPH-infested and SSB/BPH-infested plants. The four types of plants were placed in the four corners of a cage (60cm × 60 cm × 60 cm) made of 80-mesh nylon nets, respectively. Subsequently, 40 pairs of newly emerged wasps (<1 day old) were released into the cage. After 48 hr, the rice leaves with SSB eggs were collected, and the total number of SSB eggs on each plant was counted and their parasitization status was determined under a microscope two days later; the eggs turned black after being parasitized for 3 days. The experiment was replicated 12 times. The experiment was performed in a greenhouse at 27 ± 3°C and with 75 ± 10% RH and a photoperiod of 16:8 hr L:D.

### Statistical analyses

All data were checked for normality and equality of variances prior to statistical analysis. Likelihood ratio test (LR test) applied to a generalized lineal mixed model (GLMM) for overdispersion and grouped design were conducted to compare the number of eggs laid by SSB females on rice plants (Poisson error structure with log link function). Likelihood ratio test (LR test) applied to a generalized lineal model (GLM) were conducted to compare the parasitism rates of SSB eggs by *T. japonicun* (normal distribution error) with the cage treated as a random factor; the percentage data of parasitism rates were arcsin square-root transformed prior to analyses. Two-way analysis of variance (ANOVA) followed by least significant difference (LSD) test was used to compare the body weight increases of the SSB larvae on different plant treatments. The contents of JA and SA in different samples was analyzed using one-way ANOVA followed by Tukey honest significant difference (HSD) test or two-sided Student’s *t*-test. Behavioral responses of *T. japonicun* in Y-tube assays were analyzed using binomial test with an expected response of 50% for either olfactometer arm; parasitoids that did not make a choice were excluded from the analysis. Differences in volatile emission and in gene expression were analyzed by partial least squares-discriminant analysis (PLS-DA) ^25,55^ using SIMCA 14.1 software (Umetrics, Umeå, Sweden). The omics data were normalized by medians, log-transformed, and then auto-scaled (mean centered and divided by the standard deviation of each variable) using Metaboanalyst 4.0 software ^73^ before they were subjected to PLS-DA. All statistical analyses were conducted with SPSS 22.0 (IBM SPSS, Somers, NY, USA), except for the PLS-DA were performed using SIMCA 14.1 software.

## Data availability

RNA-seq raw data have been deposited to the Gene Expression Omnibus (GEO) database in the National Center for Biotechnology Information (NCBI) (note: accession number is not open yet). All other relevant data are available from the corresponding author upon reasonable request.

## Acknowledgements

The study was supported by the National Natural Science Foundation of China (31972984 and 31901896). The contribution by T.C.J.T. was supported by European Research Council Advanced Grant 788949.

## Author contributions

Y.Li conceived and directed the project. Y.Li, Q.L., X.H., and T.C.J.T. designed the study. X.H. and S.S. performed the experiments. Q.L., X.H., S.S., Y.Li, and T.C.J.T analyzed the data. Q.L., X.H., S.S., Y.P., G.Y., Y.Lou, T.C.J.T., and Y.Li wrote the manuscript. All authors have read and approved the manuscript for publication.

## Additional information

### Supplementary information

The online version contains supplementary material available.

